# Collaborative partnership model to transform bioinformatics core into a highly effective research partner and multiply the impact

**DOI:** 10.1101/2022.09.21.508957

**Authors:** R. Krishna Murthy Karuturi, Govindarajan Kunde-Ramamoorthy, Gregg TeHennepe, Joshy George, Vivek Philip

**Affiliations:** Computational Sciences, The Jackson Laboratory, USA

## Abstract

Many bioinformatics cores face a multitude of challenges. We recognized that the primary source of these challenges was the service-centric approach. So, we initiated the transformation of our bioinformatics core, Computational Sciences (CS), at the Jackson Laboratory (Jax) to be a science-centric collaborative research partner for our faculty and project stakeholders. We call our model as *collaborative partnership model*. With the effective replacement of the service model with the collaborative partnership model, CS now acts as both an effective collaborator and a co-driver of scientific research and innovation at Jax. In this paper, we describe the principles and practices we adopted to realize this transformation and present the resulting growth in the impact of CS in the research enterprise at Jax.

## Introduction

Many bioinformatics cores have been set up to serve the informatics and statistical needs of research in faculty labs, especially in wet labs. The services offered by bioinformatics cores include experimental design, analytical pipeline development, basic to advanced data analysis, software application development, and computational resource management [5].

Unlike the other scientific cores with more concrete deliverables (e.g., genotyping and tissue imaging), a bioinformatics core is often confronted with substantive changes to the deliverables as the project progresses, and as such present a significant management challenge [3-4]. For example, the design and feature requirements of a software application that seeks to provide a user-friendly analytical tool can change dramatically upon user feedback. Similarly, the direction of a data analysis project may change as the findings at each stage of the project may need unforeseen analysis to be performed.

These fundamental challenges have become formidable by the increasing breadth and complexity of biomedical projects and ever-changing analytical tools [1,3-5]. The bioinformatic activities required for these projects are wide-ranging and can include: (i) integrative analysis of increasingly complex heterogeneous omics and imaging data, (ii) algorithm development, (iii) application of machine learning (ML) and artificial intelligence (AI), (iv) process and pipeline development, (v) identification of the appropriate set of self-service analytics tools, (vi) development of novel software applications to integrate data and tools for ease of access, and (vii) deployment of tools on local high-performance computing and cloud environments for optimal performance and scaling. Effective delivery of such projects requires the core staff to work at the convergence of biology, statistics, computer science, and emerging informatics technologies [3]. In addition, bioinformatics cores face a significant compounding complexity of projects that vary dramatically in their size, nature (e.g., infrastructure, center, developmental and research projects), data availability (e.g., quality, quantity, heterogeneity, and sufficiency) [6], and project stakeholders (e.g., internal collaborators, pharma collaborators, and consortiums). These challenges become virtually unsurmountable when constrained by the transactional service-approach, which cannot scale to meet increased complexity [1-9]. Another major challenge is recruiting staff who have the skills to navigate the complex project space, and retaining them due to lack of career progression, lack of attribution of work, and appropriate rewards for their work.

To address these challenges, several recommendations were made by the heads of the bioinformatics cores: starting with Lewitter and Rebhan in 2012 [7] to the recent publications by Judith et al [6], Chang et al [3-4] and Mazumdar et al [8]. Lewitter and Rebhan’s recommendations encompass understanding the mission and organizational context of the core, staffing, staff training and support for staying connected with trends, prioritization of projects, partnerships with scientists, and conflict resolution. Though many of these recommendations are valid even today, the model of operation in many cores remain service-centric. Mazumdar et al [8] highlighted the importance of programmatic collaborations and funding allocations within their cancer center. Whereas Chang et al [3-4] highlighted the importance of appropriately rewarding and supporting staff with career development opportunities that include appropriate attribution of credit, separate career tracks, and time for research. However, they also highlighted that these challenges persist despite separate career ladders. Both groups suggest bioinformatics faculty as part of the core.

Despite the recommendations offered in the literature, the current practice largely remains a transactional service approach and the challenges remain intact even today as highlighted by Chang et al [3-4] and Julie et al [5]. The service approach typically includes a combination of practices: (i) a menu of services, (ii) a service level agreement (SLA), (iii) a statement of work (SoW), (iv) setting expectations, (v) time reporting, (vi) generating reports, and (vii) a feedback/improvement tracking system. However, these practices cannot address the above challenges. To be specific, a defined menu of services can protect the core from scope management-related issues, but it will limit the flexibility and innovation which, in turn, severely restricts the core’s ability to meet the complex, evolving bioinformatics needs of the research organization they serve. Similarly, Service Level Agreements (SLAs), meant to communicate expectations of mutual engagement with faculty, in concert with statements of work (SoWs), can help set expectations related to the scope of the work. Both mechanisms limit the flexibility needed to adapt to the natural evolution of most medium-to-complex projects and can limit the impact and collegial scientific discussion. In addition, many cores practice timekeeping and track detailed metrics of analyses, and track complaints using a feedback tracking system. Though they are useful in building trust, they have limited significance to the projects if the respective scientific questions are not answered, a new scientific discovery is not made, or a project fails to make an impact. These service-centric practices deepen separation between the core and the faculty labs which in turn may limit staff development and the ability to attract and retain talented staff who can provide scientific expertise and leadership at the bioinformatics core.

Faced with these challenges and lack of success in overcoming them with the traditional strategies using the transactional service model, ten years ago, we were motivated to replace the service-model with a new model called the *Collaborative Partnership Model* with a motto of *‘Focus on science, not service-practices’*. In the collaborative partnership model, core scientists and engineers act as scientific collaborators and take co-responsibility for the project. However, the model required us to make transformative changes in every aspect of our bioinformatics core, Computational Sciences (CS). In this paper, we describe the principles used for comprehensive organizational transformation we created to successfully transform CS into a highly effective innovative and collaborative partner for the scientists and project stakeholders.

## Principles and practices to establish a collaborative partnership model

We adopted the following principles and practices for the transformation CS to collaborative partnership model.

### 1. Establish an inspiring research-centric vision

*Bioinformatics isn’t a transactional service, it is the pursuit of scientific knowledge* [13]. Hence, we set out the purpose of CS to *co-lead innovative data science and application development that enables research and leads to important discoveries*. With this vision, we aimed to transform CS into an innovative collaborative partner unit, where our faculty find their best computational collaborators who focus on the science and shoulder significant project responsibilities.

### 2. Develop a 360-degree view of the organizational context

To execute our vision for CS, we proactively engage leaders across the institution to develop a 360-degree understanding of the tri-partite, but intertwined, organizational context: (1) the institution, (2) the faculty and the technology groups, and (3) the CS staff.

The institutional context relevant to CS is a combination of the major research & education programs, faculty recruitment strategy, intramural funding priorities, and partnerships of the institution. Furthermore, we also pay close attention to the long-term vision and mid-term strategic priorities of the institution. These provided important inputs for our agile and sustainable approach to the transformation and operation of CS.

Besides broader institutional context, beyond the ongoing projects with CS, we engage faculty and technology leaders to understand their groups’ programs, emerging directions, informatics mentoring needs of their staff, short-term informatics support needs of their projects, and most importantly, science-focused informatics expertise CS brings to the table. They all together provide sustainable collaborations and long-term embedding for CS staff in research and technology programs which have helped us in transforming our group into an effective collaborative partner.

The CS staff play a very important role in the execution of the vision and success of the core as an effective innovative and collaborative partner. Our strategy included providing exciting collaborative opportunities, support for career growth, and projects that benefits and engages CS staff. The details are described in the following principles.

Together, we have taken a proactive approach to understand the context, lay out the strategy, and execute it to the benefit of all CS stakeholders and staff. We emphasized on accomplishing the benefits for the institution (efficiency and stability of the workforce), for the faculty and technology groups (flexibility and comprehensive collaborative expertise at a phone call), and for the CS staff (stability, research and leadership opportunities).

### 3. Create a matrix structure that supports major programs and projects

We developed a bi-modal matrix structure for CS: an administrative functional structure and an agile project structure.

The administrative functional structure aligns with the major research and technology programs. At JAX, these areas included Cancer Informatics, Immuno-Informatics, Quantitative Genetics, Genome Informatics, Single Cell Genomics, AI/ML & Imaging, SQA, Systems Integration & Engineering, UI/UX & Visualization, and program management. We have appointed a leader for each program or vertical, who is responsible for planning, hiring and mentoring the associated staff to be experts and leaders in their programmatic area, and integrating with other groups within CS and across JAX.

Whereas the agile project structure is led by science and technology leads in CS and supports complex projects that require diverse skills spanning multiple verticals. For example, our PDX project requires staff who optimize genome analysis, carry out cancer data analysis, create visualizations, and automation of data analysis. A project leader draws their team from appropriate functional groups, work closely with the project sponsors, and shoulders the responsibilities for project execution. This structure is orthogonal to the administrative functional structure and offers greater agility.

Together, simultaneous operation of these orthogonal structures kept CS practically flat and integrated. As a result, we have been able to support a multitude of complex projects at JAX.

### 4. Implement programmatic embedding and matrix management of CS staff

Based on our 360-degree understanding of the Jax context and building on the above CS’s matrix structure, we embed each staff into one of the major programs at Jax so that our staff members receive programmatic mentorship and career planning from their CS functional leader while receiving project/domain mentorship from the respective collaborating faculty members and project leaders. To enhance the programmatic embedding, we adopted multi-channel flexible project intake in which the embedded staff also act as critical points of contact to the collaborators and feed that information to the CS management for further planning and resourcing of the projects. The programmatic embedding played a critical role in sustainable growth in collaborations, and attracting top talent, and development of CS staff.

### 5. Recruit researchers and engineers with leadership potential

To take up the above leadership responsibilities and proactive collaborators, we looked for candidates who, in addition to the necessary informatics skills, have a potential for and interest in leadership, a strong interest in and ability to conduct collaborative research at Jax, willingness to understand and own the scientific vision of projects, ability to manage multiple collaborations, excellent communication skills, and interest in a wide variety of informatics domains.

To attract such candidates, we revamped our career tracks to be inclusive and aspirational, with the central goal of creating a team that can offer co-leadership for research projects at Jax. Besides the analyst and statistician career tracks used in a typical bioinformatics core, we introduced the computational scientist track which was aimed to recruit collaborative researchers and project leaders. We revamped the software engineer track to scientific software engineer track to emphasize their collaborative role in science and leadership responsibility in research projects. Both tracks have five levels with an emphasis on innovation and collaboration at all levels, and special emphasis on research and project leadership at mid-to-senior levels. The progress of staff in a role depends on their demonstration of scientific/technological expertise, the leadership of complex projects/programs, impact as demonstrated by their primary and collaborative authorship on grant applications, role in project development, and scientific publications. As a result, the mid-to-senior level scientist positions emerged as quasi-faculty positions that fill important gaps between analysts and independent faculty. This clear definition of science and growth-centric career tracks, combined with the collaborative embedding, helped us attract top talent to CS which in turn reinforced our vision and drove the cultural shift in CS.

Building on our innovation and leadership-centric career tracks, we improved every step in the hiring process. We revamped our job postings to highlight collaborative research and leadership opportunities. The CV review criterion for scientists and analysts includes, besides technical skills, evaluation for strong programmatic research focus and innovative research as highlighted by both (co-)first author publications and other publications, grant awards, participation in grant authorship, research statement, and mentoring experience. A similar approach was taken for software engineers for their domain experience, as publications may not be expected of many software engineer candidates. We broadened our interview process to include CS-wide and institutional participation. The interview panel consists of teams of staff from different CS verticals as well as collaborators and leaders. The hiring managers communicate the input they seek from each team of the panel. The standard process included team-candidate interaction, seminars for scientists/analysts, and whiteboard sessions for software engineers to evaluate programmatic emphasis and problem-solving skills. Furthermore, we widened our recruitment channels with the help of our Education department, collaborators and CS staff.

### 6. Develop core staff into researchers and project leaders

To support the vision and continue the efforts in the hiring, we grow CS staff to be experts and leaders at the cutting edge of science and technologies relevant to Jax. The staff development principles are:

a. Integration of staff into programs by identifying projects and programmatic collaborations of interest to them.
b. 360-degree mentoring on all fronts of their role in our core: collaborative interactions, project management, communication, and science & technology leadership. We use both performance dialogue as well as monthly discussions between staff and managers to accomplish it.
c. Develop T-shaped expertise which develops deep expertise in one area of bioinformatics such as cancer informatics or visualization while having a broad understanding of the skills required for complex bio-medical projects. Such a skill profile helps the staff to understand the complexity of the project, have seamless conversations with their team members, see their work in the larger context, and make them successful team players and key contributors to complex bio-medical projects. To this end, we require staff to take up informatics courses outside their core area of work e.g., all staff are required to take genetics & genomics courses, software engineers are encouraged to take courses in analytics, and biologists to take up computer science courses.
d. Provide research and leadership opportunities. We actively explore co-leadership opportunities for staff in the grant applications, identifying suitable intra-/extra-mural grant application opportunities our staff can apply for, and supporting project development in the emerging technologies, helping them to identify long-term collaborations, and offering supervisory opportunities.
e. Encourage staff to explore a multitude of career opportunities (sci & tech leadership, administrative functional leadership, and project leadership) available at CS and Jax. We actively understand their aspirations and identify suitable career followed by making necessary investments for each member of CS.
f. Conduct annual staff engagement surveys, retreats. The surveys help us get input from our staff to make necessary adjustments to our management and mentoring methodology, and the retreats engage them in the vision and trajectory of the CS. The staff engagement is also driven by staff participation and leadership in the new initiatives. In addition, the flexible work models and engagement with Diversity-Equity-Inclusion exercises further support staff growth.
g. Regular training for managers in management and communication which are essential in creating a conducive environment for open discussion and hence growth opportunities for staff.

### 7. Establish partnership with IT and other relevant departments across the institution

Information Technology (IT) is an important component of our everyday work and successful delivery of our projects. To this end, we have established partnership with IT group by working together in identifying, testing and deployment of relevant technologies and IT practices. Besides, the partnership has been realized via regular open communication, joint retreats, and working for mutual success. Our partnerships extended beyod IT, with all key relevant groups and departments: Education, Grants, and Human Resources. We prefer to be partners rather than customers with these critical groups as we value mutual engagement on opportunities, aspirations and limitations, and finding creative ways of identifying paths forward. The Education department channels trainees to CS, helps CS staff to participate in developing the educational strategy and educational deliverables, and provide relevant training for CS staff. Our partnership with the Grants office helps identify suitable funding opportunities, track grant applications that may require CS participation and establish collaborations in the project’s inception, and train CS staff in successful grant applications. For example, we got a feature added to the information system that helps us track the grant application proposals so that we can proactively touch base with the program leaders at the project’s earliest stage to plan for the appropriate staff to be embedded in the program. It also helped us plan for the recruitment of the appropriate talent and the re-training of current staff. Our partnership with Human Resources helps us develop good career tracks and grade them to be in line with the job market, enforcing uniform use of these tracks across the organization, developing manager training programs, and developing CS into an organization.

### 8. Cultivate a culture of collaborative team data science and establish a computational community

Instituting a collaborative culture is another critical element for the success of our vision. We used several strategies to accomplish it.

First, besides programmatic embedding, we co-located CS staff amidst the faculty labs. With the embedding, the CS staff deliver formal training programs to develop citizen data scientists, mentor trainees and junior staff, provide resource coordination among faculty labs and the core, ensure end-to-end engagement in projects. In addition, we encouraged collaborations by developing a repository of analysis pipelines which enabled not only efficiency, but also uniformity in practice and collaborative culture.

Second, the team data science approach plays a key role in establishing collaborative culture. Another dimension of modern bio-medical projects is the complexity stemming from their multi-disciplinary nature. To address this problem, like many data science organizations in the industry, we decided to develop unicorn teams that could deliver every skill and coordination essential for such projects. These teams are led by one of the experienced staff from CS or a faculty lab. The choice of the team lead isn’t dependent on the title or job type, but rather dependent on the interest in the leadership and ability to understand the multi-disciplinary members of the team.

Third, our emphasis on the integration of computational staff from faculty labs and CS is another key driver to establish a collaborative culture. Besides informal mentoring for faculty lab members by CS staff, a formal computational community was formed. The community is engaged with working groups along with bi-annual retreats. The community working groups are identified and led by a team of computational staff from CS, IT, faculty labs, and technology groups. The computational community is governed by a committee of staff from CS, IT, and faculty labs and services group. Program managers from CS and faculty labs collectively organize this committee and the community.

### 9. Develop and practice Research Project management

As a result of programmatic embedding of CS staff and our collaborative team data science approach, the relationship and communication problems were greatly reduced but not eliminated. These issues persisted due to the complex, data-intensive, and exploratory nature of bio-medical projects. To address this dimension of the problem we introduced Research Project Management (RPM) methodology. Major principles of our RPM methodology include: (1) knowing the scientific vision of the project by using the grant application or the project proposal as a reference point, (2) knowing the audience of the project, (3) understanding the multi-disciplinary matrix management, (4) flexible scope, schedule, and resource management, (5) monitoring and communication of progress, (6) joint team leadership by a research project manager and a sci/tech lead, and (7) engaging all stake holders and the team of the project. The engaged include faculty, supervisors of the team members, and partners of the project from across Jax. We used a two-pronged approach to deliver RPM: we train CS staff in the RPM methodologies to manage low to medium complex projects without adding administrative overhead, and developed a dedicated RPM group to manage complex long-term projects.

RPM has proven to be central to ensuring the consistent delivery of scientific results, improved planning and communication, and significantly increasing collaborator satisfaction and trust in CS. RPM has been supported by project management software. Overall, the RPM approach effectively established a open, transparent, and efficient workflow at JAX.

### 10. Measure and track the transformation and impact of CS

During the transformation, we realized that typical core-styled metrics (recovery, budget, and project wait times) do not drive the vision, justify the necessary investments, and identify the changes needed. Based on the premise that, “what we measure is what we optimize”, we defined and tracked several metrics to drive the collaborative science-centric culture. These included: (i) number of total and lead-authored publications of CS staff, (ii) number of grant applications supported and contributed to as primary authors, (iii) number of pipelines deployed and updated, (iv) number of trainees in the faculty labs mentored by our staff, (v) number of education modules delivered, (vi) number of citizen data scientists developed, and (vii) number of complex projects managed. Overall, the metrics captured the *scientific, collaborative, mentoring* and *leadership* performance of CS staff and CS. Whereas, the success of our managers is measured by the progress of the vision laid out, collaborator satisfaction, staff development, attrition rates, group productivity, and leadership. Today, these metrics provide a realistic and comprehensive view of the contributions of our core and are highly appreciated at JAX as measures of performance of CS leading to continued investment in CS by our collaborators and the senior management.

### 11. Plan for growth with multi-modal delivery

The transformation has been much more impactful leading to the explosion of demand for CS participation. To handle rapid growth in demand, we have taken number of steps: reorganization of CS as its size and demand surpasses the capacity of the existing structure, growing current staff into leadership roles, working with the JAX management on adjusting the funding as appropriate, and managing the expectations of the research enterprise.

Furthermore, during the transformation, we faced the challenge of balancing the effort allocation for projects of varying complexity and impact. For example, transactional bioinformatics requests are typically small and can be conducted in a day. Whereas the long-term collaborative projects require programmatic embedding and allocation of significant effort over a long period. Hence, we developed different modes of delivery depending on their complexity and impact: automation of analyses by developing and democratizing workflows, offering consultation for faculty lab staff who have significant computational skills but require expert input, created helpdesk to address non-programmatic requests that can be completed within a day, mentoring scientists in faculty labs to conduct low complexity analysis, and co-lead complex multi-disciplinary projects. The decisions are taken by the CS management in consultation with faculty and the team members.

An important aspect of growth planning is the communication of the impact of CS on research and the (co-)leadership provided by CS staff to the research enterprise. We introduced a quarterly newsletter that communicates metrics and the associated details. The newsletter is sent to the whole research enterprise.

## Results

The transformation of CS resulted in numerous benefits to JAX. JAX benefits with the stability and efficiency of the scarce and valuable human capital, and effective complex team data science for the research enterprise; flexible, efficient, complex project resourcing along with bioinformatics partnership for faculty; and an enriching computational ecosystem that offers professional mentoring, career planning, and most importantly independence, flexibility, and professional development to the computational staff resulting in low staff attrition rates at CS compared to similar groups. In specific, the following areas of growth highlight the impact of our model.

### Multifold growth in Funding and Staffing

Our total funding from intramural and extramural grants increased by >20x from 2012 to 2021 with CS effort on these grants increased from 2FTE to ∼40FTE. During the same period, the staffing has grown only by 4-fold. Such tremendous and efficient growth was possible as our vision and its execution using the above principles attracted top talent to CS which in turn attracted more collaborations and co-leadership opportunities, led to the virtuous cycle. Staffing in the key areas increased by more than 6-fold. We forecast this growth to continue as more faculty invest in and collaborate with CS.

### Structure

The growth has been supported by two rounds of the significant revision of the structure of CS. We identified emerging technologies and programmatic areas at Jax and aligned the new structure with it. As a result, the number of thematic groups increased from 3 to 12 with a matrix structure co-led by managers and sci/tech leads.

### Analytics evolved to support complex heterogeneous data and algorithm development

Capabilities, complexity, and productivity on the analytics front have seen huge growth. We developed and maintained 35+ workflows for processing bulk and single-cell sequencing of mouse and human omics data. The workflow repository includes optimized workflows for patient-derived xenografts (PDX) data. We have tremendously increased our capabilities of data analysis: from RNA-seq/WGS/Exome-seq data analysis for quantification/SNP calls in 2012 to a variety of bulk sequencing data, single-cell sequencing data, spatial transcriptomics data, and imaging data for diverse inferences e.g., SNPs, SVs, alternative splicing, cell type prediction, etc in 2021-22. These are all in addition to successfully collaborating with faculty on integrating this heterogeneous data to infer the biology of development and disease.

### Software Applications delivered are multi-disciplinary and outward-facing

We had one external-facing software application developed and maintained in FY12. It was a database application, with no analytics embedded. By FY21, we had multiple software applications that host the data with analysis tools embedded. A significant number of them are external facing e.g., Mouse Phenome Database (MPD) [14], Gene Weaver [15,16], Clinical Knowledgebase (CKB) [17], and Human Phenome Ontology (HPO) database [18]. In addition, several image and video processing applications have been developed for use within the technology groups at Jax. These applications include machine learning and data processing components too.

### Projects are multi-disciplinary, and their complexity increased multi-fold

During this period, we have taken multi-disciplinary projects of multi-fold complexity and are successfully supported by cross-functional teams with a team data science approach. The projects we carry out today include data from a multitude of technologies and software application development. They include integrative analysis of heterogeneous data to algorithm development. We support this complex environment with a team of four research project managers who also play a significant role in the staff integration and efficient management of our matrix structure.

### Leadership in research and development

CS scientists acted as PIs, Co-PIs, or collaborators for >15 grant applications a year and co-authored more than 120 articles in the recent 5 years with primary authorship in >40% of them. In addition, our CS staff regularly initiate grant applications in response to the internal and external funding opportunity calls, be corresponding authors on several publications, and deliver talks and participate in panel discussions at national and international conferences and workshops.

We have highlighted these growth areas along with the associated fold-change over FY12-21/22 period, in the table below.

| **Growth Area** | **Metric** | **Fold Change**<br>**2012-21** |
| --- | --- | --- |
| Funding | (\$ ) Recovery | 24x |
|  | FTE on grants | 20x |
| Staffing | Total Staff | >4x |
|  | PhD Staff | 8x |
|  | ML&AI Staff | 25x |
|  | Senior Staff | 6x |
|  | Managers/Directors | 4x |
| Structure | Major Informatics themes | 4x |
|  | Computational Community | New |
|  | Community Working groups | New & Active |
| Analytics | Seq Libraries/year | 90x |
|  | Analytics Pipelines | >35x |
|  | Technologies dealt | 9x |
|  | Projects with integrative analysis | >40x |
| Software Applications | Integrative applications (Analytics + Software) | >5x |
| Projects | Team Projects | 20x |
|  | Multi-disciplinary projects | 10x |
|  | Project Managers | >3x |
|  | Faculty supported | ~2x |
|  | Services supported | 7x |
| (Co-) Leadership | Grant Applications | >20x |
|  | Publications – Total | >2x |
|  | Publications – primary authorship | 2x |
|  | Invited presentations | 15x |

## Discussion

We accomplished the goal of transforming CS into a scientific organization that partners and closely collaborates with faculty labs, while also delivering its function as a bioinformatics core. Although our model differs from the original intent of bioinformatics cores, it overcomes the challenges inherent in a typical transactional service approach. Based on our experience, a bioinformatics core will be more successful if it effectively integrates with faculty labs and become indispensable research partner. This requires the ability to offer the expert, domain-specific collaboration that is essential for facilitating faculty in addressing their scientific questions.

Our model offers additional opportunities for the core as well as computational staff across the research enterprise. It will benefit the whole research enterprise in several areas that tremendously improve effectiveness and efficiency of the bioinformatics enterprise. This includes retention of knowledge, setting up common processes and pipelines, mentoring trainees, and cross-pollination of programs which can offer more opportunities to ask even bigger questions. We are at the inception of such a distributed enterprise at Jax with many working groups actively engaging each other.

However, the journey of transformation wasn’t without challenges. They include getting organizational buy-in, a cultural transformation towards collaboration and engagement, and establishing collaborations with partner departments. These challenges require constant transformative guidance by the leadership of the core, mentoring of the core staff to take up the collaborative leadership role, and steadily getting buy-in from the whole scientific enterprise.

Despite these challenges, this is an important journey for a core to take on and a very important direction for the respective scientific enterprises to invest in. However, it is important to note that paying attention to all principles outlined in this paper and in Judith et al [6] are important to achieve such transformation.

Overall, for CS, we proceeded from one state to a nearest impactful new state in a step-by-step manner. For example, we transformed the service model to a collaborative model in the 1^st^ phase. In the next phase, we transformed it to a co-leadership model. Now we initiated transformation to a pioneering innovation model.

## Acknowledgements

We thank Prof. Ed Liu (former CEO and President of JAX) who set us on this important journey, Dr. Ken Fasman (the SVP of Research at Jax) who guided us through this complex journey to the current state, Dr. Lon Cardon (CEO … President of JAX) for continued support, and the JAX’s-senior management team for their unwavering support for the transformation. We thank Profs. Charles Lee … Nadia Rosenthal (Scientific Directors of Jax), Dr. Madeleine Braun (CPI of Jax), Dr. Alan Sawyer (Sr Director of Scientific Services), Profs. Jeff Chuang … Greg Carter (faculty partners of CS), the faculty … heads of all sci/tech departments who actively supported and guided the model as it was being developed, and the CS leads … staff who relentlessly pursued the model and helped realize the vision. We also thank Dr. Auro Nair (former EVP of JMCRS at Jax), Ms. Val Scott (former Sr Director of Scientific Services), and Profs. Gary Churchill … Carol Bult (former faculty partners of CS) for their support and guidance in the beginning of the journey. The work has been partly supported by the JAX Cancer Center Computational Sciences Shared Service (P30 CA034196).

